# Efficient and sensitive profiling of RNA-protein interactions using TLC-CLIP

**DOI:** 10.1101/2022.04.13.488168

**Authors:** Christina Ernst, Julien Duc, Didier Trono

## Abstract

RNA-binding proteins are instrumental for post-transcriptional gene regulation, yet transcriptome-wide methods to profile RNA-protein interactions remain technically challenging. We present an improved library preparation strategy for crosslinking and immunoprecipitation (CLIP) that involves tailing and ligation of cDNA molecules (TLC) for increased sensitivity and efficiency. TLC-CLIP eliminates time-consuming purifications, reduces sample loss and minimises experimental steps, allowing precise profiling of RNA-protein interactions from limited starting material at nucleotide resolution.

---

Protein-centric approaches to study RNA-protein interactions mainly rely on cross-linking and immunoprecipitation (CLIP) of RNA-binding proteins (RBPs)^1–5^. Over the years, several variations of this technique emerged, most prominently iCLIP^6,7^ along with derivations such as enhanced CLIP (eCLIP)^8^, infrared CLIP (irCLIP)^9^ and more recent improvements including iCLIP2^10^ and improved iCLIP (iiCLIP)^11^. These techniques enable the mapping of RNA binding sites at nucleotide resolution, and while individual steps differ between protocols, they follow the same overall strategy: cells are cross-linked with UVC light followed by lysis and partial RNA digestion before or after immunoprecipitation of the RBP of interest. Copurified RNA is then 3’adapter ligated prior to SDS polyacrylamide gel electrophoresis (SDS-PAGE) and transferred onto nitrocellulose from where RNA is liberated, purified and reverse transcribed into cDNA prior to second adapter ligation and amplification to generate sequencing-compatible libraries.

Major bottlenecks during the library preparation include extensive purification steps and suboptimal enzymatic reactions that lead to sample loss, low complexity libraries and the requirement for large amounts of starting material.

Here we present an improved library preparation strategy that offers time-efficient and sensitive generation of sequencing libraries from low input material.

Our approach relies on **<u>t</u>**ailing and **<u>l</u>**igation of **<u>c</u>**DNA molecules (TLC), which enhances enzymatic reactions and employs a fully bead-based, single-tube strategy that minimises sample loss prior to amplification (Fig. 1a). This novel design makes purification of RNA-protein complexes via SDS-PAGE optional, thus providing the potential for a fully automated CLIP workflow for high-throughput settings.

**Fig. 1.**
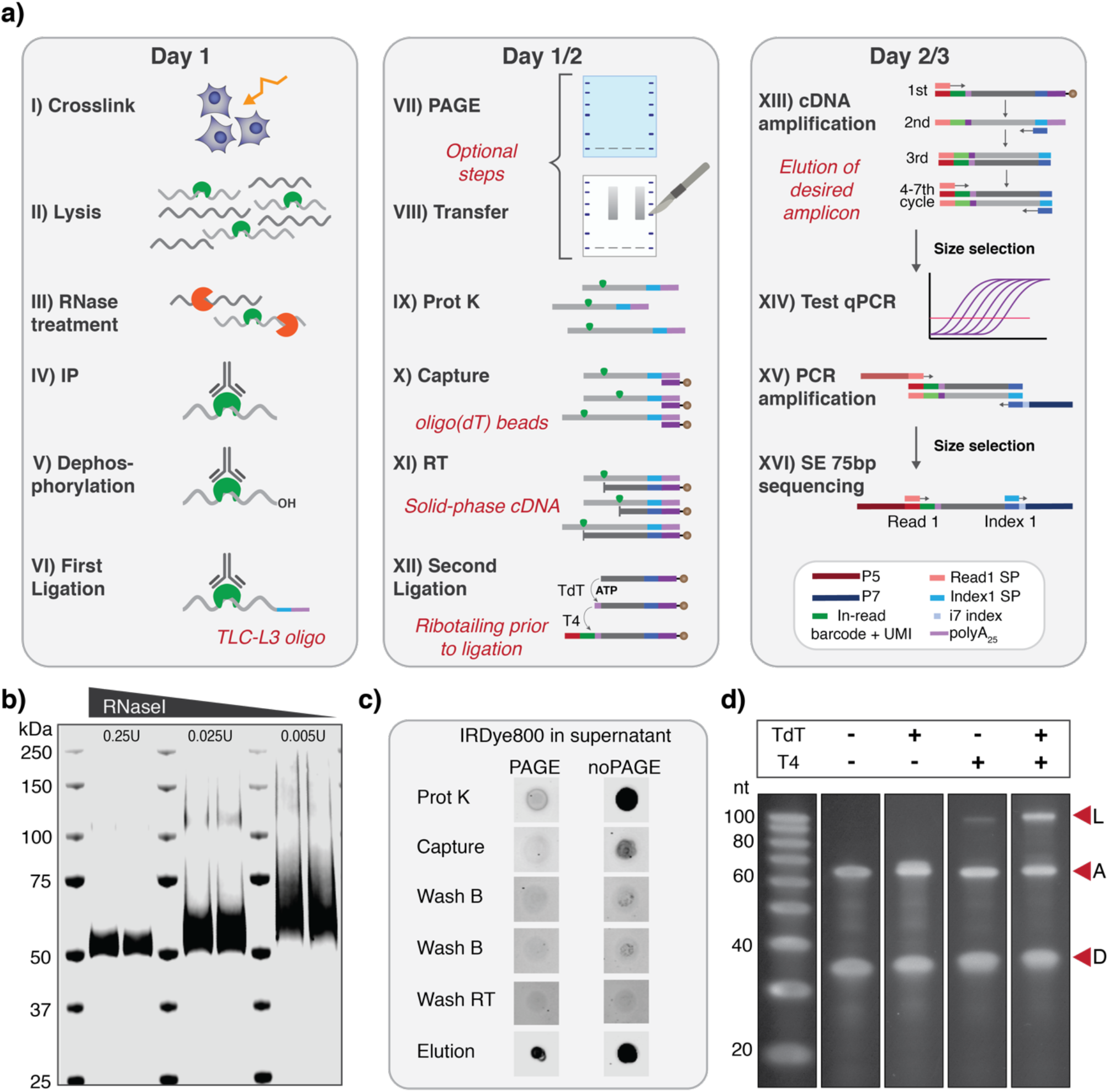
TLC library preparation enables efficient and time-effective generation of CLIP libraries. **a** | Schematic overview of TLC-CLIP procedure with major changes highlighted in red italics. **b** | Visualisation of adapter-ligated RNA on LI-COR Odyssey Clx Imager after membrane transfer for hnRNPA1 samples treated with different RNase concentrations. **c** | Capture and elution of adapter-ligated RNA throughout steps IX - XI of library preparation visualised via dotblotting of supernatants on nitrocellulose. **d** | Addition of Terminal Deoxynucleotidyl Transferase (TdT) in ligation reaction results in strongly increased ligation efficiency. D indicates donor molecule; A indicates acceptor molecule; and L the ligation product.

The majority of current CLIP-related protocols purify RNA-protein complexes via SDS-PAGE followed by transfer onto nitrocellulose, at which point crosslinked RNA can be visualised to control IP efficiency and RNase treatment conditions using radioactive isotope labelling^3,4^. TLC-CLIP employs an infrared-dye conjugated adapter that was adopted from the irCLIP protocol^9^, which avoids radioactive labelling and allows easy visualisation of adapter-ligated RNA following transfer, as well as during subsequent library preparation steps (Fig. 1b,c & Supplementary Fig. 1a,b).

The TLC-L3 adapter further contains a poly(A) stretch, which offers several advantages throughout the library preparation. First, adapter-ligated RNA molecules can be captured using oligo(dT) beads, which are inert to high concentrations of proteinase K and denaturing agents, thereby eliminating the need for time-consuming RNA precipitations that are prone to sample loss especially at low concentrations. Instead, oligo(dT) capture is highly efficient, occurs within minutes, and allows stringent washes to remove traces of proteinase K prior to subsequent enzymatic reactions (Fig. 1c & Supplementary Fig. 1c). Furthermore, oligo(dT) beads are used to prime the reverse transcription reaction, resulting in cDNA molecules covalently linked to magnetic beads, enabling efficient separation from adapter-ligated RNA via heat denaturation (Fig. 1c).

Solid-phase cDNA can then directly serve as acceptor molecule in the second adapter ligation without any additional purification procedures, thus minimising sample loss. To date, the second adapter ligation presents a major bottleneck during library preparation; a problem which is not unique to CLIP, but also presents a challenge in other protocols where random priming of second strand synthesis is not favourable or feasible^6^. Current CLIP protocols tackle this step either through circularisation of cDNA molecules^6,9,11^, or direct single-stranded (ss)DNA ligation^8,10^, but both strategies are known to be inefficient, resulting in the permanent loss of molecules that fail to ligate.

We have improved the efficiency of the ssDNA ligation approach by incorporating Terminal Deoxynucleotidyl Transferase (TdT) in the ATP-containing ligation mix, which results in the addition of a short ribo-tail to the 3’ end of cDNA molecules, greatly increasing the affinity of T4 RNA ligase (Fig. 1d & Supplementary Fig. 2a,b)^12,13^. TdTs are highly processive in the presence of deoxynucleotide triphosphates (dNTPs), but self-terminate after incorporating only a few nucleotide triphosphates (NTPs) (Supplementary Fig. 2c,d)^14^. This results in an over-representation of T nucleotides at only the first few base positions of TLC-CLIP reads, which are removed during processing (Supplementary Fig. 3).

Ligated cDNA molecules are eluted off the beads and simultaneously pre-amplified using PCR with short primers (Supplementary Fig. 4), followed by size-selection to enrich for molecules with insert sizes larger than 20 nucleotides. Sequencing-ready libraries are then generated via PCR amplification through the addition of P5 and P7 sequences harbouring i7 indeces, followed by size-selection of fragments larger than 160bp (Supplementary Fig. 5). TLC-CLIP libraries are compatible with single-end, two-colour chemistry sequencing protocols, with the first 15 nucleotides of the reads corresponding to a 9-nt unique molecular identifier (UMI) for deduplication that is split around a 6-nt barcode for greater multiplexing capacity (Methods).

The improved library preparation strategy greatly increases sensitivity and lowers input requirements, which we demonstrated by generating high-quality libraries for four different RBPs from only 50.000 cells. We then tailored a streamlined analysis pipeline^15^ to the specificities of the TLC-CLIP workflow (Figure 2a) and overall obtain a larger fraction of ‘usable reads’ compared to public CLIP datasets (Supplementary Fig. 6a-d & Table 1). Furthermore, TLC-CLIP libraries display a longer read length and a higher fraction of reads carrying deletions indicating frequent read-through instead of truncation at the crosslinking site (Supplementary Fig. 7). This is in accord with the use of Superscript IV during reverse transcription, which frequently causes crosslinking-induced mutations (CIMS)^16^, resulting in insertions and deletions at similar or higher rates compared to HITS-CLIP (Supplementary Fig. 7e,f). However, unlike in HITS-CLIP protocols, truncated reads are also retained during TLC-CLIP resulting in much greater yield of usable reads, thus drastically lowering sequencing requirements and cost compared to HITS-CLIP (Supplementary Fig. 6d).

**Fig. 2.**
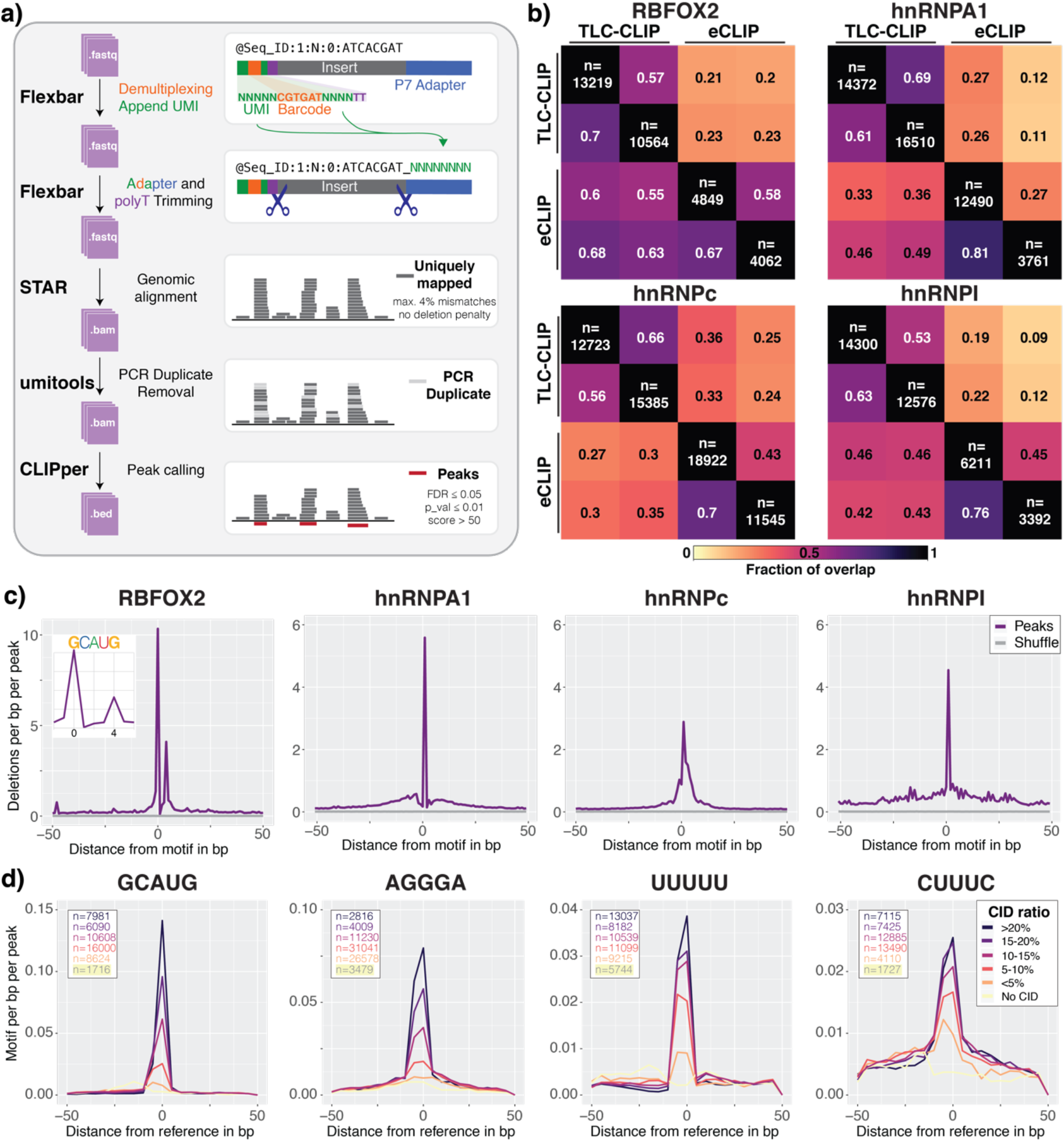
TLC-CLIP generates high-quality data with increased resolution from crosslinking-induced deletions. **a** | Schematic representation of the data processing pipeline for TLC-CLIP libraries. **b** | Fraction of overlap at the peak level between TLC-CLIP and eCLIP libraries for four different RNA-binding proteins, requiring a minimum overlap of 25% between peaks. Peaks are restricted to genes with similar expression level between 293T and HepG2 cells (n = 1507) and individual number of peaks per replicate is annotated in heatmap. **c** | Density plot showing enrichment of deletions across peaks which were centred on the corresponding consensus motif for each RBP. For control regions, peaks were shuffled across target gene bodies of a given RBP. **d** | Density plot of consensus motif across peaks subset based on their crosslink induced deletion (CID) ratio (n_del_/n_reads_) at the reference point, which either presents the deletion maximum or apex region for peaks without deletions.

Given these characteristics, we opted for the peak calling algorithm CLIPper, which does not exclusively rely on read start positions to determine cross-linking events^8,17^, and can therefore be applied to different protocols, allowing a direct comparison between TLC-CLIP and public CLIP datasets. TLC-CLIP libraries show a high level of correlation between peaks comparing individual replicates (R^2^=0.5-0.78, Pearson correlation), and up to 50%overlap with eCLIP peaks despite differences in the cell types that were profiled (Supplementary Fig. 8a,b). Restricting the comparison to genes with similar expression levels between 293T and HepG2 cells, or comparing TLC-CLIP and easyCLIP^18^, which were both performed in 293T cells, increases the overlap to up to 72%, which is similar to technical variation observed between replicates, given the stochastic nature of RNA binding (Fig. 2b and Supplementary Fig. 8c). Furthermore, de novo motif discovery on TLC-CLIP peaks recapitulates previously reported motifs with high precision and shows strong enrichment around the peak apex, which represents the most likely region of crosslink defined by CLIPper (Supplementary Fig. 9).

The precision of TLC-CLIP can be further enhanced by incorporating the positional information of crosslinking induced deletions (CIDs), which are highly correlated between replicates at the singlenucleotide level (R^2^=0.47-0.62, Pearson correlation) (Supplementary Fig. 10a). Deletions are strongly enriched at RBP binding motifs and show high precision, identifying the two guanines of the canonical RBFOX binding motif ‘GCAUG’ as crosslinking sites^19^ (Fig. 2c). Thus, centring peaks on the position with the highest number of deletions increases precision, as well as motif enrichment, which is strongest in peaks with a higher ratio of crosslinking-induced deletions, highlighting the benefit of using CIDs to improve the specificity of TLC-CLIP data (Fig. 2d and Supplementary Fig. 10b).

This is of particular interest when applying TLC-CLIP without PAGE purification, which enables a 2-day fully automatable workflow and further lowers the input requirements down to 500 cells. Libraries generated without PAGE purification recapitulate the binding behaviour of a given RBP, capturing the position-dependent enrichment in relation to splice sites and intronic Alu elements for RBFOX2 and hnRNPc, respectively (Fig. 3). We observe up to 67% overlap at the peak-level between libraries generated with or without PAGE purification, but with lower motif enrichment in peaks specific to libraries omitting PAGE, indicating a higher level of contaminating background sequences, as expected when removing an additional purification step (Supplementary Fig. 11a, b). Lower motif enrichment is accompanied by lower CID ratios, demonstrating the utility of CIDs as an additional quality filter to discern true binding sites from copurifying, non-crosslinked fragments in samples with higher background signal, which can also result from sub-optimal RNase conditions (Supplementary Fig. 11c-e). Annotation of peaks with lower CID ratios shows an increase of coding sequences (CDS), which are a common background contamination also frequently detected in eCLIP libraries, demonstrating that filtering based on CIDs provides increased specificity to remove background signal without the need to generate matched input samples (Supplementary Fig. 12)^11^.

**Figure 3:**
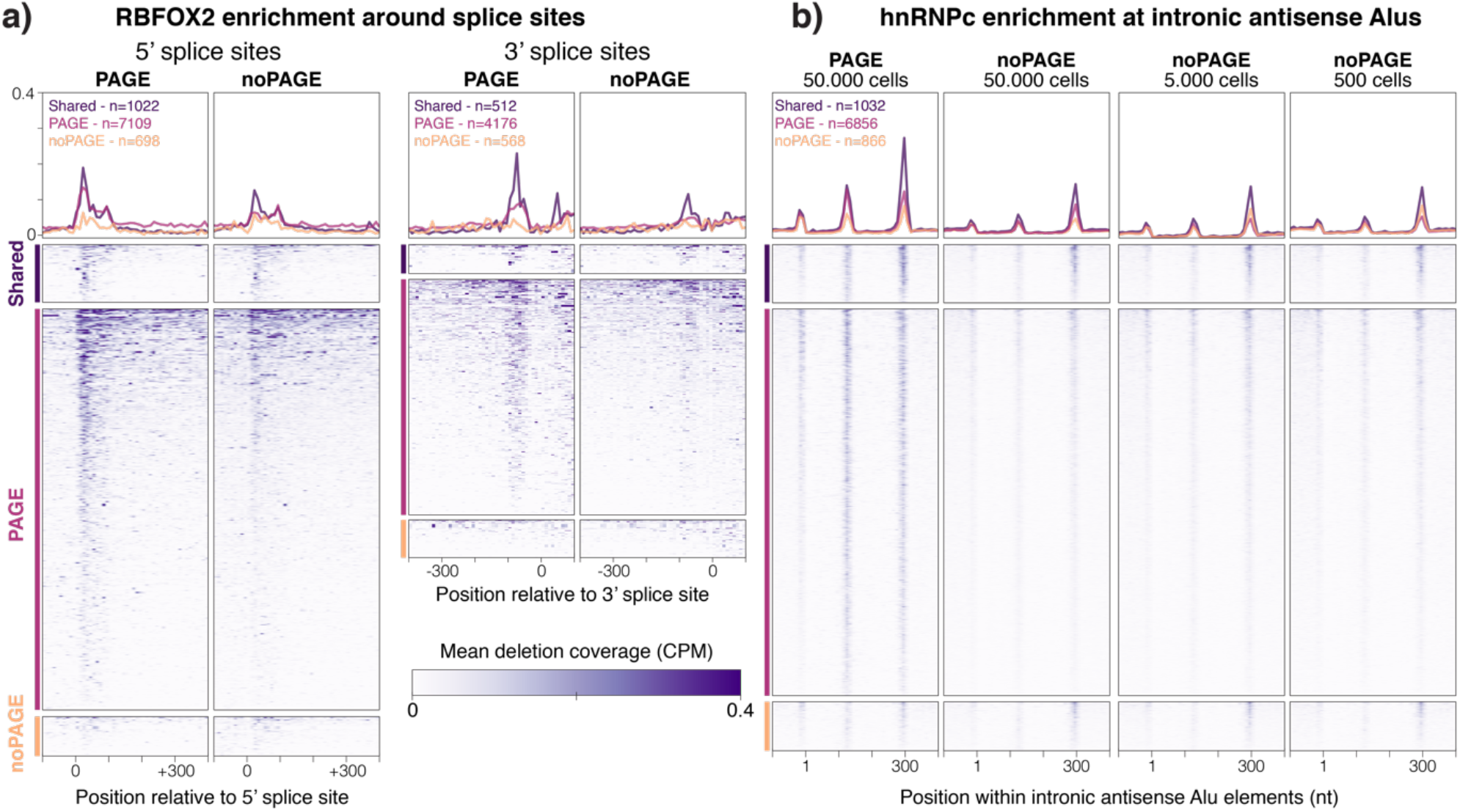
Crosslinking-induced deletions enable precise position-dependent binding visualisation. **a** | Deletion coverage of RBFOX2 TLC-CLIP libraries prepared with and without PAGE purification in relation to 5’ and 3’ splice sites across peaks with a CID ratio > 10 (CID10) in both PAGE and noPAGE libraries (shared), or peaks for which this threshold was met in only one of the experimental conditions (PAGE or noPAGE). **b** | Deletion coverage across intronic Alu elements in antisense orientation to the host gene for hnRNPc TLC-CLIP libraries. Heatmaps are divided in Alu elements overlapping CID10 peaks in both PAGE and noPAGE libraries (shared) or Alu elements which overlapped a CID10 in only one condition. Libraries omitting PAGE purification for hnRNPc were prepared from 50.000 cells as well as serial dilutions of the same input material down to 500 cells.

Taken together, the streamlined library preparation protocol of TLC-CLIP drastically reduces both time and cost of CLIP experiments, while generating high quality RBP binding profiles from low input material. The larger number of crosslinking induced deletions further improves both the precision and specificity of TLC-CLIP libraries, by increasing the nucleotide resolution of crosslinking sites and distinguishing true binding sites from co-purifying, non-crosslinked fragments. Furthermore, by eliminating the need for PAGE purification, input requirements can be further reduced with high quality data obtained from as little as 500 cells, presenting a fully bead-based, single-tube library preparation strategy amenable to automation for high-throughput settings.

## Methods

### TLC-CLIP adapter generation

All adapters and oligos used throughout the protocol were ordered from Integrated DNA Technologies (IDT) and information regarding sequences, scale and purification can be found in Supplementary Table 2.

The TLC-L3 oligo for the first adapter ligation was synthesized at 250 nmole scale, carrying a 5’ phosphorylation and 3’ IRDye^®^ 800CW (NHS Ester) (v3) modification and was purified using RNase-free HPLC with a total yield of 21.1 nmoles. Pre-adenylation was performed on 5 nmoles using the 5’ DNA Adenylation Kit (NEB, E2610L) as follows: 50 μl of 100 μM L3 adapter were set up with 25 μl 10X 5’ DNA Adenylation Reaction Buffer, 25 μl 1 mM ATP and 50 μl Mth RNA Ligase (1nmol) in a total volume of 200 μl. Reaction was incubated at 65°C for 2 hours followed by inactivation at 85°C for 10 minutes, during which it turns cloudy. Reaction was then cleaned up using the Nucleotide Removal Kit (Qiagen, Cat #28304) as follows: 200 μl were mixed with 4.8 ml of PNI buffer, distributed over 10 columns and spun down at 6000 rpm for 30 seconds. Columns were washed once in 750 μl PE buffer, spun for 1 min at 6000rpm, followed by an empty spin at full speed before transferring columns to a new collection tube. 50 μl H2O were added per column and incubated at RT for 2 minutes before centrifugation at 6000 rpm for 1 minute. Eluates were combined with an approximate final concentration of 10 μM and 1 μM working stocks were prepared and frozen at - 20C. Aliquots can be freeze-thawed at least 20 times without any detectable loss in activity.

### Cell culture and generation of CLIP lysates

Adherent 293T cells (ATCC^®^ CRL-1573^™^) were grown to ~80% confluency in Dulbecco’s Modified Eagle Medium (Gibco, #41966-029) supplemented with 10% FCS (Sigma-Aldrich, #F9665-500ML, Lot #19A124) and 1% Penicillin-Streptomycin-L-Glutamine (MED30-009-CI). Cells were rinsed in ice-cold PBS and crosslinked on ice with 254 nm UV-C light at 0.3 J/cm^2^ in a CL-3000 Ultraviolet Crosslinker (UVPA849-95-0615-02). Cells were collected into PBS by scraping, counted and desired cell number was aliquoted and spun down. Cell pellets were resuspended in iCLIP Lysis buffer (50 mM Tris-HCl pH 7.4, 100 mM NaCl, 1% Igepal CA-630, 0.1% SDS, 0.5% sodium deoxycholate) using 50 μl per 50.000 cells. Lysates were incubated on ice for 5 minutes followed by sonication for 5-10 seconds at 0.5 seconds ON and 0.5 seconds OFF at 10% amplitude using a tip sonicator (Branson LPe 40:0.50:4T). Protein concentration was measured using the Pierce^™^ Rapid Gold BCA Protein Assay Kit (Thermo Scientific, A53225) and lysates were either processed directly or stored at −80°C.

### RNase treatment, immunoprecipitation and first adapter ligation

Protein-G beads (Thermo Fisher, 10004D) were washed twice in 1ml iCLIP Lysis buffer and resuspended in 100 μl per condition (100 μl of protein-G beads bind 20-30 μg of antibody - scale accordingly). Per IP, 1 μg of antibody against hnRNPc (Santa Cruz Biotechnology, sc-32308), RBM9 (Bethyl Laboratories, A300-864A), hnRNPA1 (4B10) (Santa Cruz Biotechnology, sc-32301), or hnRNPI (Santa Cruz Biotechnology, sc-56701) were added, and antibody-bead mixture was incubated at room temperature (RT) for 30-60 minutes on a rotating wheel.

Meanwhile cell lysates were treated with different RNase concentrations using 0.25U, 0.025U and 0.005U of RNase I (Thermo Fisher, EN0602) for high, medium and low conditions. RNase dilution was added to cell lysates together with 2ul Turbo DNase (Thermo Fisher, #AM2238) and lysates were incubated at 37°C for exactly 3 minutes at 1100rpm, followed by 3 minutes on ice. Cell lysates were spun down for 10 minutes at 4°C at full speed and supernatant was transferred to a new tube.

Antibody-bead mixture was washed twice in iCLIP lysis buffer to remove unbound antibody and RNase-treated cell lysates were added alongside cOmplete EDTA-free Protease Inhibitor Cocktail (Merck, #11836170001) and incubated for 2 hours at 4°C on a rotating wheel. After IP, beads were washed twice in 200 μl High Salt Buffer (50 mM Tris-HCl pH 7.4, 1 M NaCl, 1 mM EDTA, 1% Igepal CA-630, 0.1% SDS, 0.5% sodium-deoxycholate), with the second wash at 4°C for 3 minutes on a rotating wheel, followed by two washes in 200 μl PNK Wash Buffer (20 mM Tris-HCl, pH 7.4, 10 mM MgCl2, 0.2% Tween-20).

Dephosphorylation of 3’ ends was performed in 20 μl of PNK reaction for 30 minutes at 37°C (70 mM Tris-HCl, pH 6.5, 10 mM MgCl2, 1 mM DTT, 10U SUPERaseIN RNase Inhibitor (ThermoFisher, #AM2696), 5U T4 Polynucleotide Kinase (NEB, #M0201L). Beads were washed twice in PNK Wash Buffer and resuspended in 20 μl of ligation mix for overnight incubation at 16°C and 1200 rpm (50 mM Tris-HCl, pH 7.8, 10 mM MgCl2, 1mM DTT, 10U SUPERaseIN RNase Inhibitor, 10U T4 RNA Ligase (NEB, #M0204), 1 μl of 1 μM L3 adapter and 20% PEG400 (Sigma-Aldrich, #91893)).

### TLC-CLIP library preparation with PAGE purification

Following the first adapter ligation, beads were washed twice in 200 μl High Salt Buffer, twice in 200 μl PNK Wash buffer and then resuspended in 20 μl 1X LDS sample buffer (Thermo Fisher, #NP0008) containing 5% betamercaptoethanol (Sigma-Aldrich, #M6250). Samples were denatured for 1 minute at 70°C and RNA-protein complexes were resolved on NuPAGE 4-12% Bis-Tris Gels (Thermo Fisher, #WG1402A) at 180V for 1 hour. Transfer was performed onto nitrocellulose (BioRad, #1620115) in 1X NuPAGE transfer buffer (Thermo Fisher, #NP00061) with 10% methanol at 30V for 2 hours at RT.

Nitrocellulose membranes were scanned on Odyssey^®^ CLx Infrared Imager (LI-COR, 9141) with 169 μm resolution to visualise RNA localisation and then placed on filter paper soaked in PBS. Regions of interest were cut out from nitrocellulose membrane corresponding to ~20-100 kDa above the molecular weight of the RBP of interest due to the ligation of L3 adapter (~15.9 kDa) and associated RNA (with 70nt of RNA averaging ~20kDa). Nitrocellulose pieces were placed in LoBind Eppendorf tubes and 200 μl Proteinase K buffer (100mM Tris-HCl, pH 7.4, 50 mM LiCl, 1 mM EDTA, 0.2% LiDS) containing 200 μg Proteinase K (Thermo Fisher, #AM2546) were added and incubated at 50°C for 45 minutes at 800rpm.

Meanwhile, 10 μl of Oligo(dT)25 Dynabeads™ (Thermo Fisher, #61005) per sample were washed in 1 ml of oligo(dT) Binding Buffer (20 mM Tris-HCl, pH 7.4, 1 M LiCl, 2 mM EDTA) and resuspended in 50 μl of oligo(dT) Binding Buffer per sample. Following Proteinase K treatment, supernatant was transferred to fresh tubes containing 50 μl of oligo(dT) beads and incubated for 10 minutes at RT on a rotating wheel. Following RNA capture, beads were washed twice in 125 μl oligo(dT) Wash Buffer (10 mM Tris-HCl, pH 7.4, 150 mM LiCl, 0.1 mM EDTA) and once in 20 μl 1X First-Strand Buffer (50 mM Tris-HCl, pH 8.3, 75 mM KCl, 3 mM MgCl2). Beads were resuspended in 10 μl of Reverse Transcription Mix (1X First-Strand Buffer, 0.5 mM dNTPs, 1 mM DTT, 6U SUPERase IN RNase Inhibitor, 20U SuperScript™ IV Reverse Transcriptase (Thermo Fisher, #18090050) and incubated for 15 minutes at 50°C followed by 10 minutes at 80°C heating up to 96°C. Samples were vortexed for 30 seconds at 96°C and then immediately placed on a magnet on ice. Supernatant containing adapter-ligated RNA was removed and efficiency of elution can be confirmed by dot-blotting on nitrocellulose membrane.

Solid-phase cDNA was washed once in 60 μl oligo(dT) Wash Buffer and once in 20 μl 1X T4 RNA Ligase Buffer (50 mM Tris-HCl, 10 mM MgCl2, 1mM DTT, pH 7.5). Beads were resuspended in 5 μl of 5’ Adapter mix (2 μl 10X T4 RNA Ligase Buffer, 2 μl of 10 μM L## oligo (see Supplementary Table 2), 1 μl 100% DMSO), incubated at 75°C for 2 minutes then immediately placed on ice. 4 μl of Ligation Mix (5 mM ATP, 7U Terminal Deoxynucleotidyl Transferase (TdT) (2230B, Takara), 15 U T4 RNA Ligase High Concentration (M0437, NEB)) were added as well as 10 μl 50% PEG8000 and reaction was mixed by pipetting up- and down until beads are resuspended. Reaction was incubated at 37°C for 30 minutes, then cooled down to room temperature. 30 U of T4 RNA Ligase were added, the reaction mixed by pipetting and incubated at RT overnight with occasional vortexing for 15 seconds at 2000 rpm every two minutes.

Following overnight incubation, ligation reaction was removed and beads washed in 100 μl oligo(dT) Wash Buffer and 20 μl 1X Phusion HF Buffer (Thermo Fisher, #F518L). Beads were resuspended in 25 μl cDNA amplification mix (1X Phusion HF PCR Master Mix (NEB, #M0531L) and 0.5uM P5 and P7 short primer mix (see Supplementary Table 2)) and amplification was performed with the following programme: 30 seconds at 98°C, 7 cycles of 10 seconds at 98°C, 30 seconds at 65°C and 30 seconds at 72°C followed by final extension at 72°C for 3 minutes. Meanwhile, 2μl of oligo(dT) beads per sample were washed once in 1 ml oligo(dT) Binding buffer and resuspended in 5 μl per sample. After cDNA amplification, 5 μl of oligo(dT) beads were added and incubated at RT for 5 minutes on a rotating wheel to capture unwanted amplification byproducts (see Supplementary Fig. 4). Samples were placed on magnet and supernatant containing amplified cDNA was transferred to a fresh tube.

Size-selection of cDNA was performed using ProNEX^®^ Size-Selective Purification System (Promega, #NG2002) with a ratio of 2.8X to enrich for cDNA inserts of at least 20 nucleotides in length (>80bp). Library yield was then estimated by amplifying 1 μl of purified cDNA via qPCR using the full length P5 and P7 index primers and 2-3 cycles were subtracted from the obtained Ct value for final library amplification. Following PCR amplification, libraries were size-selected again using the ProNEX^®^ Size-Selective Purification System, with a ratio of 1.8X to select fragments larger than 165bp. Quality control was performed using the Agilent High Sensitivity DNA Kit (Agilent, #5067-4626) and libraries were quantified using the KAPA Library Quantification Kit (Roche, #KK4824).

### TLC-CLIP library preparation without PAGE purification

When omitting PAGE purification, the first adapter ligation was performed for 75 minutes at 25°C. Beads were washed as described above and either directly resuspended in Proteinase K reaction or in 20 μl of RecJ adapter removal reaction (1X NEB Buffer 2 (NEB, #B7002S), 25U 5’ Deadenylase (NEB, #M0331S), 30U RecJ endonuclease (NEB, #M0264S), 10U SuperaseIN and 20% PEG-400) and incubated at 37C for 30 minutes prior to Proteinase K treatment. Samples were then placed on magnet, and supernatant was transferred to fresh tubes containing oligo(dT) beads, with the remaining library preparation performed as described above.

### Sequencing

TLC-CLIP libraries were sequenced on an Illumina NextSeq500 using the High Output Kit v.2.5 for 75 cycles, using Illumina protocol #15048776. 5% PhiX were added to final library pools for increased complexity and sequencing run was performed with custom configuration, running 86 cycles for Read 1 and 6 index cycles.

### Mock ligations and denaturing polyacrylamide gel electrophoresis

Efficiency of second adapter ligation was tested in mock ligations using TLC-CLIP L01 as donor molecule and i7-3 as acceptor. 2 μl 10X T4 RNA Ligase Buffer, 1μl 10μM TLC-CLIP L01 oligo, 1μl 10μM i7-3 oligo and 1μl DMSO were mixed and incubated at 75°C for 2 minutes. Reaction was placed on ice and 4 μl of Ligation mix containing 0.2μl 0.1M ATP, 0.5 μl TdT and 0.5 μl T4 RNA Ligase High Concentration were added followed by addition of PEG8000 to the indicated percentage. Ligation was incubated for 30 minutes at 37°C then cooled down to 16°C. Half the reaction was removed after 60 minutes at 16°C, the remaining reaction was incubated overnight.

1μl of Ligation reaction was mixed with 1μl Gel Loading Buffer II (Thermo Fisher, #AM8546G) and denatured at 72°C for 3 minutes. Samples were separated on 10% TBE-Urea gels (Thermo Fisher, #EC68752BOC) and stained with 1X SYBR^®^ Gold (Thermo Fisher, #S11494) for 10 minutes in TBE buffer.

### Data analysis

#### Demultiplexing and Trimming with Flexbar

Sequencing data was demultiplexed by i7 index reads using bcl2fastq without any read trimming. Further demultiplexing by in-read 5’ barcodes and trimming of adapter sequences was performed using Flexbar v.3.5.0 (https://github.com/seqan/flexbar)^20^ in a two-step approach. In the first step, reads are demultiplexed by in-read barcodes allowing no mismatches, and UMIs are moved into the read header. Barcode sequences (see Supplementary Table 2) including the UMI designated by the wildcard character ‘N’ are provided in fasta format, with the arguments “b barcodes.fasta --barcode-trim-end LTAIL --barcode-error-rate 0 --umi-tags”. In the second step, any adapter contamination at the 3’ end of the reads is removed allowing an error rate of 0.1 with the following arguments “--adapter-seq ‘AGATCGGAAGAGCACACGTCTGAACTCCAGTCACNNNNNNATCTCGTATGCCGTCTTCT GCTTG’ --adapter-trim-end RIGHT --adaptererror-rate 0.1 --adapter-min-overlap 1”. In addition, potential T-stretches at the 5’ end that are the result of ribotailing during ligation are removed by trimming ‘T homopolymers of 1-2 nucleotide length (Supplementary Fig. 3) using “--htrim-left T --htrim-max-length 2 --htrim-min-length 1” and reads shorter than 18 nucleotides post trimming are discarded by “--min-read-length 18”.

#### STAR Alignment

Flexbar-trimmed reads were aligned against hg19 using STAR v.2.7.3a (https://github.com/alexdobin/STAR)^21^ with the following parameters, to keep only uniquely mapping reads, removing the penalty for opening deletions and insertions and fully extending the 5-prime end of reads to preserve the end of cDNA molecules: “--outFilterMultimapNmax 1 --scoreDelOpen 0 --scoreInsOpen 0 -- alignEndsType Extend5pOfRead1”. To retain UMI in read header during STAR alignment, any space in header needs to be removed prior to mapping.

#### Deduplication of Reads

Aligned reads were deduplicated based on unique molecular identifiers using UMI-tools v.1.0.1 (https://github.com/CGATOxford/UMI-tools)^22^.

The dedup command was used with the parameters “--extract-umi-method read_id --method unique --spliced-is-unique” to group reads with the same mapping position and identical UMI, while treating reads starting at the same position as unique if one is spliced and the other is not.

#### Peak Calling

Enriched regions were identified using the peak calling algorithm CLIPper v.2.0.0 (https://github.com/YeoLab/clipper)^8,17^ with default settings and a p-value cutoff of 0.01 “--poisson-cutoff 0.01”.

#### Multiqc and usable reads

General quality metrics of libraries were assessed using FastQC v0.11.7 (https://github.com/s-andrews/FastQC) and QC data were collated using multiqc v.1.9 (https://github.com/ewels/MultiQC)^23^ to extract information from combined log files to plot usable read fractions.

#### Deletions

Individual nucleotide positions of crosslink-induced deletions within TLC-CLIP reads were extracted using the htseq-clip tool (https://github.com/EMBL-Hentze-group/htseq-clip)^24^ with the following parameters: “htseq-clip extract --mate 1 --site d”.

#### Filtering of peaks

CLIPper peaks were filtered by removing ENCODE blacklisted regions from eCLIP libraries as well as peaks obtained from TLC-CLIP libraries skipping the ligation step as well as IgG controls for either Rabbit or Mouse IgG depending on RBP. An additional score filter was applied by requiring −10 log(pval) to be larger than 50 for any downstream analysis. Consensus peaks between replicates were obtained using bedtools intersect^25^ requiring a minimum overlap of 25% between peaks.

#### Correlation plots

For correlation plots peaks or deletion positions of individual replicates were concatenated and coverage was calculated using bedtools multicov^25^. Count data was normalised using the cpm function from edgeR^26^ against total library size and log2 transformed. Point density plots were generated using the geom_pointdensity package available on Bioconductor and correlation coefficient was calculated using Pearson correlation.

#### Pairwise comparison at peak level

Fraction of overlap between filtered peaks for individual replicates of either TLC-CLIP, eCLIP or easyCLIP was calculated using the intervene pairwise intersection module (https://intervene.readthedocs.io/en/latest/index.html)^27^ requiring a minimum of 25% overlap between peaks. For comparison between TLC-CLIP and eCLIP in HepG2 cells shown in Fig. 2b, peaks were restricted to genes with stable gene expression between the two cell lines, as defined by differential gene expression analysis on total RNA-seq data for 293T and HepG2 cells. Stable genes were defined as having an absolute fold change lower than 1.1 and an expression higher than 5 log2 CPM.

#### De *novo* motif discovery

*De novo* motif discovery was performed using Homer^28^ v4.10 on peaks centred on either the apex region obtained from CLIPper or after centring peaks on the position with the highest deletion count. findMotifsGenome.pl was used with the parameters

“-oligo -basic -rna -len5 -S10 -size given” where peak size is a 50-nucleotide window around the apex or with parameter “size 50” for peaks centred on deletions.

#### Density plot for deletions and motif enrichment

Motif densities were calculated using the annotatePeaks.pl function from homer on consensus motifs generated with the seq2profile.pl function allowing 0 mismatches. For density plots in Supplementary Fig. 9, motif density was calculated for apex-centred peaks from TLC-CLIP and eCLIP libraries using “--size 1000 -hist 5 - norevopp”. For hnRNPC “-rm 10” was specified to remove occurrences of the same motif within 10 nucleotides to avoid artificial amplification of motif enrichment through longer U-stretches, for hnRNPI the density of pyrimidine stretches was plotted, based on a ‘YYYYY’ stretch.

For plotting the deletion density in Fig. 2c, TLC-CLIP peaks were centred onto the consensus motif, with motif files being generated using seq2profile.pl. Tag directories for deletions were generated using the homer makeTagDirectory function on the bed file obtained from htseq-count. peakSizeEstimate needs to be changed to 1 in tagInfo.txt file to avoid extension of deletion tags and preserve nucleotide resolution. Deletion enrichment was obtained using the annotatePeaks.pl with “-hist 1 -size 100” across deletion-centred peaks as well as peaks shuffled across the set of target genes bound by a given RBP. For RBPs recognising palindromic sequences such as ‘AGGGA’ or ‘CUUUC’ for hnRNPA1 or hnRNPI respectively, the exact position of the crosslinking site cannot be determined during alignment if the deletion falls within the homopolymer stretch. By default, STAR will position the deletion at the first base of the ambiguous sequence based on the DNA sequence, without awareness of the strand orientation of the gene, resulting in an artificial shift of the deletion position between genes on the forward or reverse strand. To remove this artifact, deletion positions for genes on the reverse strand were shifted by two nucleotides for hnRNPA1 and hnRNPI prior to visualisation.

#### Deletion-centred analysis

Peaks were centred on the maximum deletion position and coverage of this nucleotide position was calculated using bedtools multicov to calculate the CID ratio, indicating the proportion of reads at a given position that carry a deletion. Motif density across peaks with different CID ratios was calculated using annotatePeaks.pl with “-size 100 - hist 5 -norevopp”.

For visualising the percentage of peaks carrying motifs according to CID ratio, findMotifsGenome.pl was used with “-find motif - size 50 -norevopp”. Peak annotation across different transcriptomic and genomic features was performed using annotatePeaks.pl.

#### Deletion Visualisation

Splice site annotation from homer for hg19 was used and intersected with deletion-centred peaks for RBFOX2, with a CID ratio larger than 10 from PAGE or noPAGE libraries, yielding splice sites that were either shared between experimental conditions or specific to either PAGE or noPAGE libraries. The same was done for hnRNPc peaks and intronic antisense Alu sequences that were extracted from RepeatMasker^29^.

Deletion positions from htseq-clip were merged across all replicates and converted to bam files using bedtools bedtobam. Bigwig files were then generated using deeptools function bamCoverage with a binsize of 1, normalising for total deletion count (CPM). Heatmaps and coverage profiles were generated using the createMatrix and plotHeatmap function from deeptools^30^.

#### Data visualisation

Downstream data analysis and visualisation was performed in R (v 4.1.0) using the tidyverse package^31^.

## Data Availability

Sequencing data have been deposited on Gene Expression Omnibus under accession number GSE200432 and are currently under curation.

## Acknowledgements

We thank Brian Zarnegar and Paul Khavari for providing reagents and guidance on initial irCLIP experiments. We thank the Gene Expression Core Facility (GECF) at EFPL for technical support. This research was supported by the European Research Council (D.T. - KRABnKAP, #268721; Transpos-X, #694658), the Swiss National Science Foundation (C.E. - PZ00P3_202048; D.T. - 310030_152879 and 310030B_173337), the Human Frontiers Science Programme (C.E. - LT000147/2019), the European Molecular Biology Organisation (C.E. - ALTF 516-2019) and the EPFL-Stanford Exchange programme.

## Contributions

C.E. designed and performed experiments, performed computational analysis, interpreted the data and wrote the manuscript. J.D. performed computational analysis. D.T. oversaw the work and reviewed the manuscript.

## Figure Captions

**Supplementary Fig. 1.**
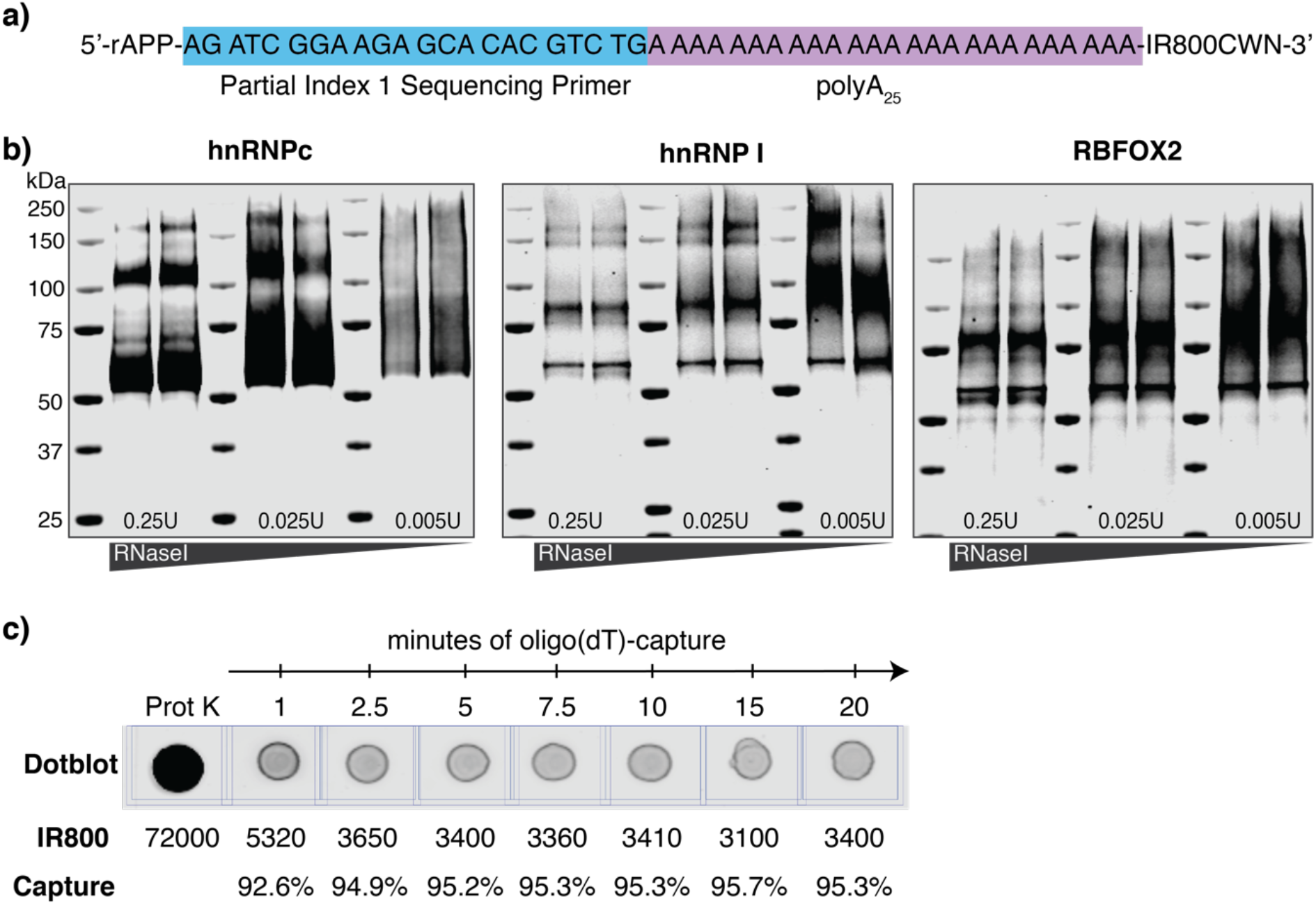
TLC-L3 oligo enables easy visualisation and efficient capture on oligo(dT) Dynabeads^®^. **a** | Sequence and design of TLC-L3 adapter consisting of partial Illumina Index 1 Sequencing Primer, a 25-nucleotide long polyA stretch and covalently coupled IR800 dye at the 3’ end for visualisation and chain termination. **b** | Visualisation of adapter-ligated RNA following transfer onto nitrocellulose membrane, displaying different RNase conditions for hnRNPc, hnRNPI and RBFOX2. **c** | Dotplot depicting timecourse of TLC-L3 capture on oligo(dT)_25_ Dynabeads after Proteinase K reaction.

**Supplementary Fig. 2.**
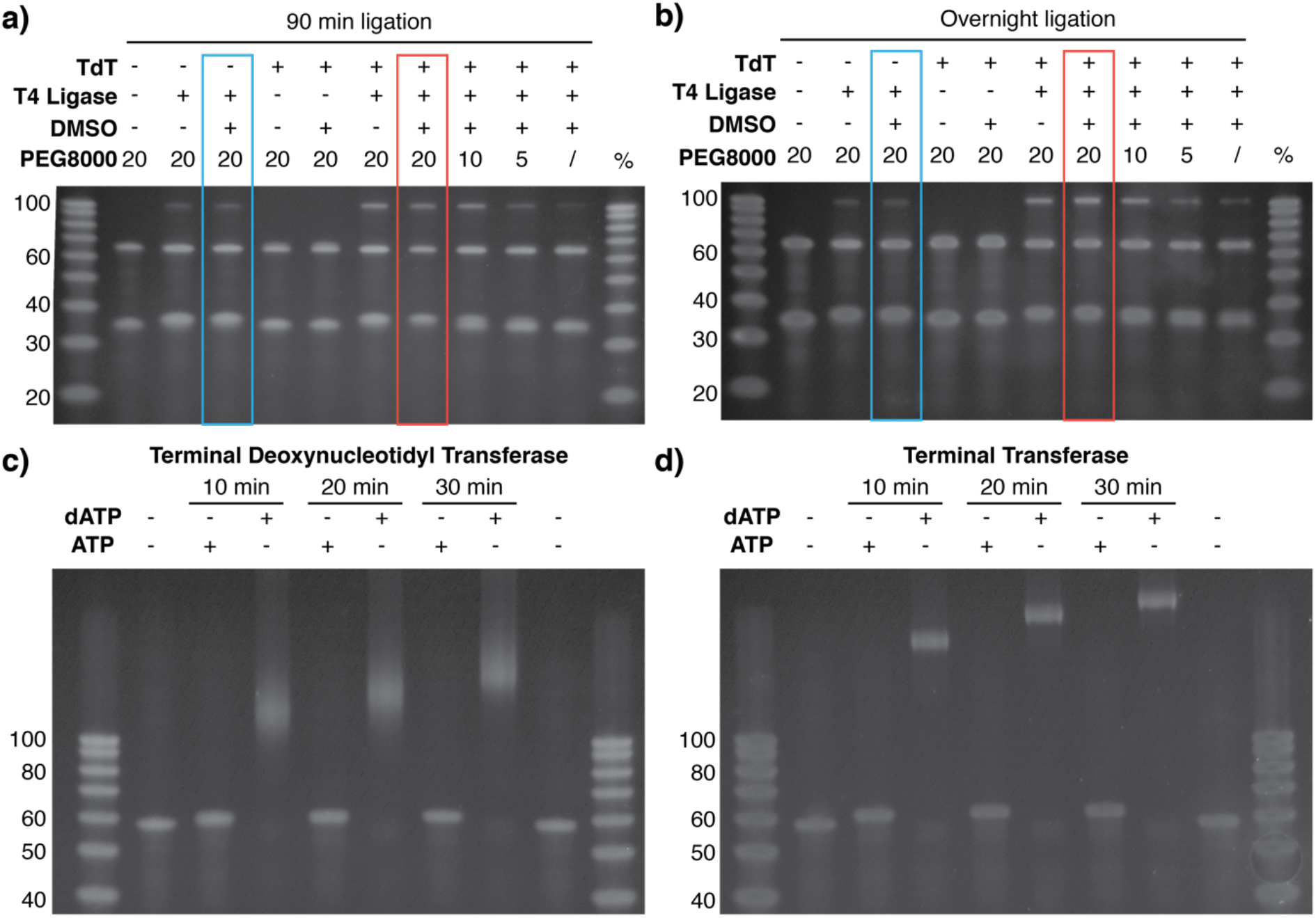
TdT improves ligation efficiency via the addition of short ribotails. **a**-**b** | Denaturing 10% TBE-Urea PAGE of mock ligation, testing different experimental conditions after (a) 90 minutes or (b) overnight ligation. eCLIP and iCLIP2 ligation conditions are highlighted by blue rectangles, TLC-CLIP ligation conditions are highlighted by red rectangles. **c-d** | Denaturing 10% TBE-Urea PAGE of tailing reaction performed with (c) Terminal Deoxynucleotidyl Transferase (TdT) or (d) Terminal Transferase in the presence of either ATP or dATP and termination by heat denaturation after 10, 20 or 30 minutes.

**Supplementary Fig. 3.**
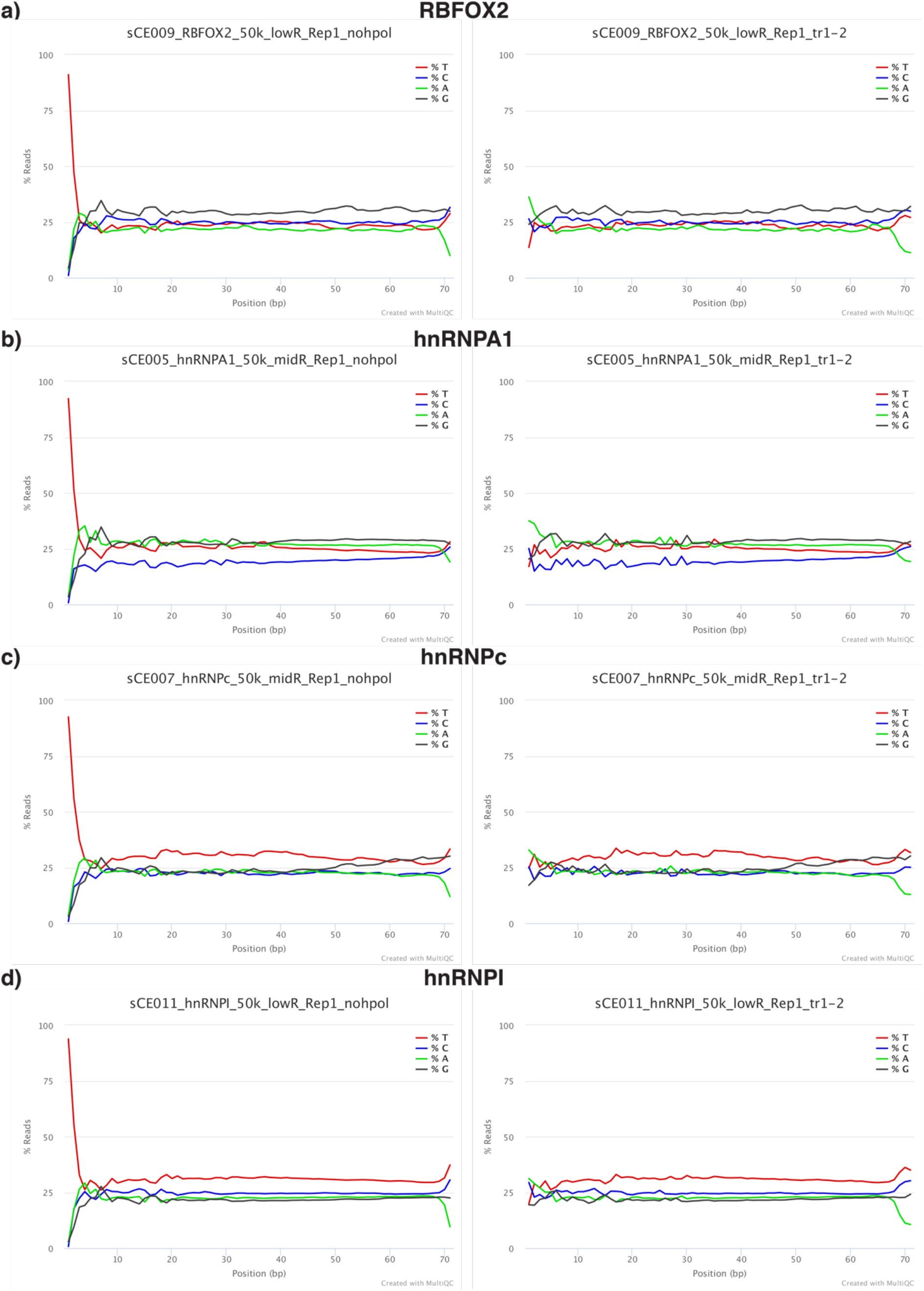
Homopolymer trimming eliminates sequence bias introduced by ribotailing during second adapter ligation. **a-d** Per base sequence content before (left) and after (right) homopolymer trimming of 1-2 T bases for different RBPs.

**Supplementary Fig. 4.**
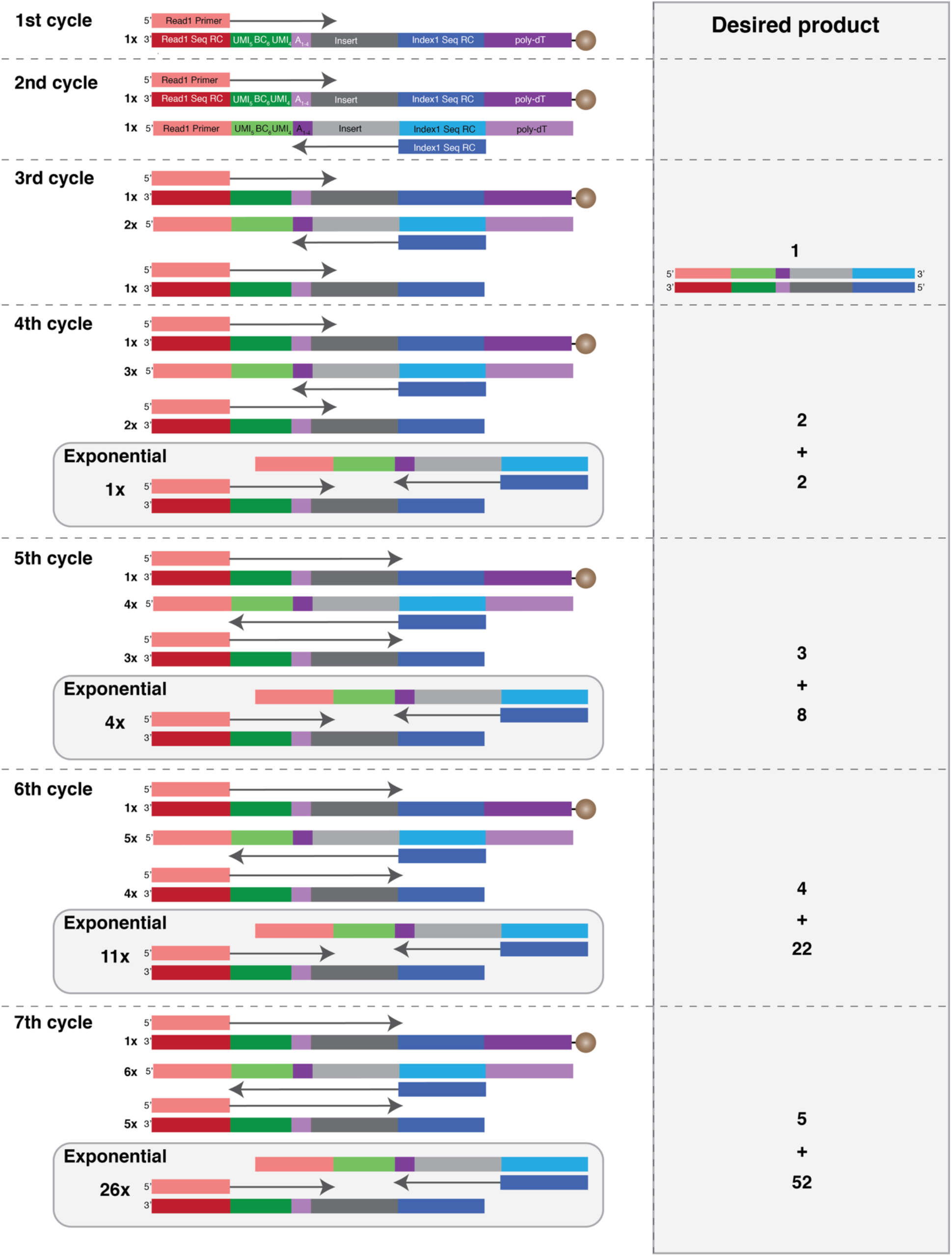
Pre-amplification scheme of cDNA. Schematic representation of cDNA pre-amplification displays the generation and elution of desired amplicons in the 3^rd^ cycle via linear amplification. Combination of linear and exponential amplification over the following 4 cycles results in 57-fold amplification. For libraries omitting PAGE purification, 6 cycles equivalent to a 26-fold amplification are sufficient.

**Supplementary Fig. 5.**
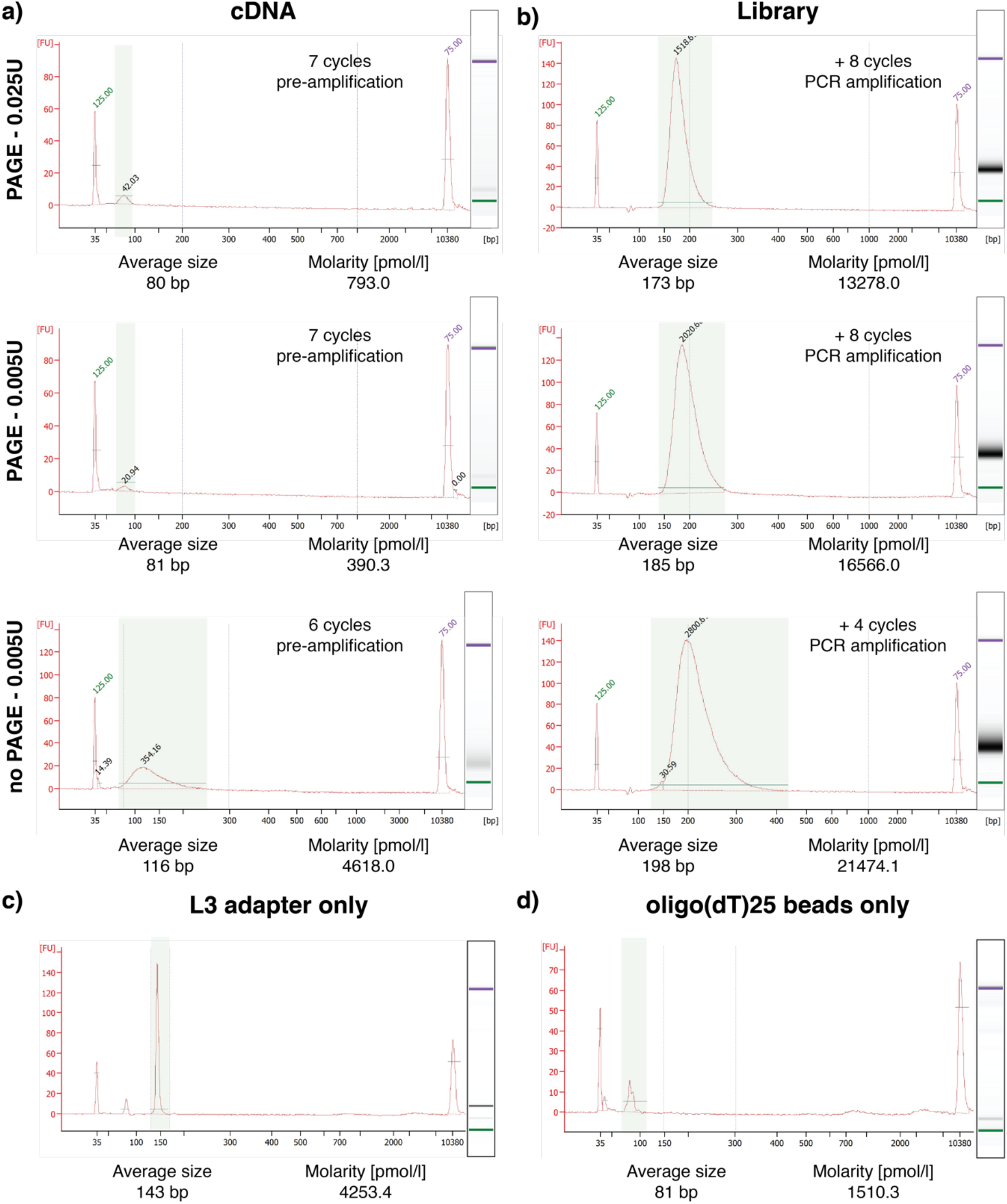
Quality control of cDNA and sequencing libraries. **a-b** | Representative examples of High Sensitivity DNA Bioanalyser traces for (a) pre-amplified cDNA and (b) sequencing-ready libraries showing PAGE-purified libraries generated with different RNase concentrations and a library generated without PAGE purification for hnRNPA1. **c-d** | High Sensitivity DNA Bioanalyser traces of control conditions, such as library preparation with (c) TLC-L3 adapter or (d) oligo(dT) beads only. Average size of area marked in green as well as molarity are stated below the graphs.

**Supplementary Fig. 6.**
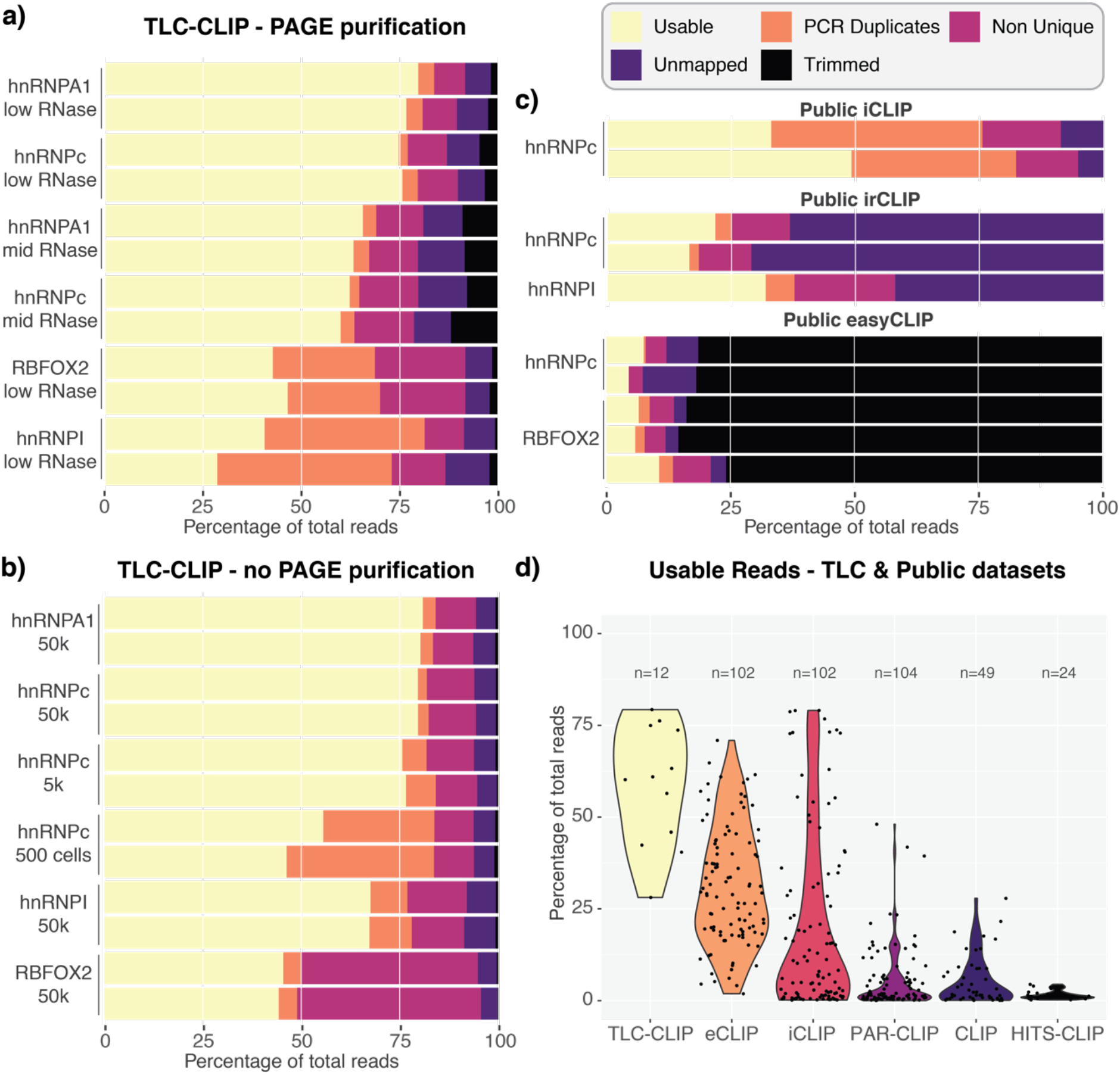
TLC-CLIP libraries retain a larger fraction of usable reads compared to other protocols. **a-b** | Percentage of total reads for different TLC-CLIP libraries are plotted based on five different categories: uniquely mapping and PCR-deduplicated (Usable), uniquely mapping PCR duplicates (PCR Duplicates), non-uniquely mapping (Non Unique), not mappable due to short size or other reasons (Unmapped) or lost during trimming due to short size (Trimmed). **c** | Same representation of read fractions is shown for publicly available iCLIP (E-MTAB-1371), irCLIP (GSE78832), and easyCLIP (GSE131210) datasets that were reanalysed for this manuscript. **d** | Percentage of usable reads out of total read fraction is displayed for different publicly available CLIP protocols and PAGE-purified TLC-CLIP libraries. Information on usable read fraction for public datasets was obtained from van Nostrand et al. (2016) and further annotated according to different experimental protocols.

**Supplementary Fig. 7.**
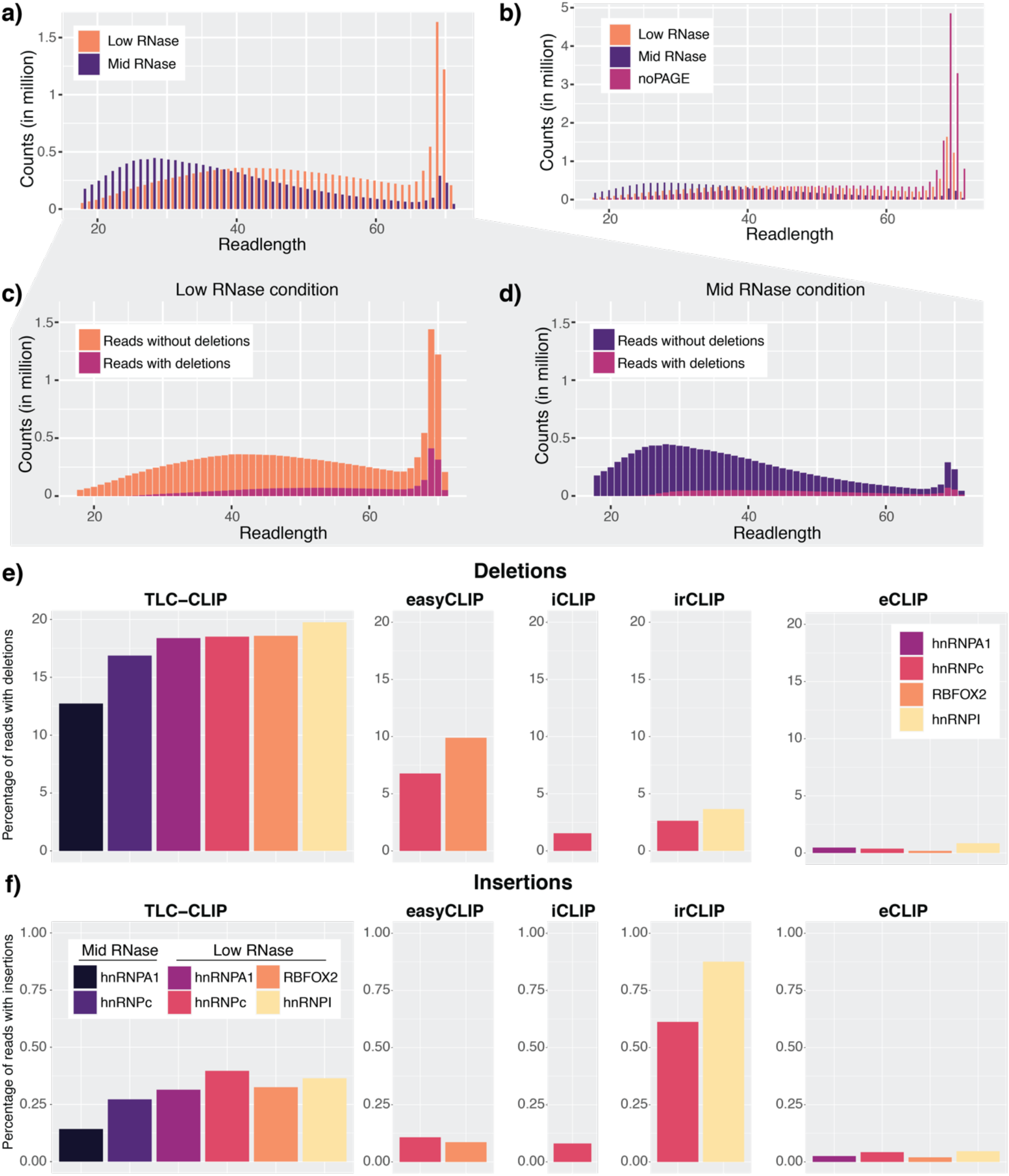
TLC-CLIP libraries display longer reads and a greater proportion of reads with crosslinked induced deletions. **a-b** | Read length distributions for hnRNPA1 comparing (a) different RNase conditions in PAGE purified libraries and (b) omission of PAGE purification. **c-d** | Read length distribution for reads with and without deletions for (c) low and (d) medium RNase conditions. **e** | Percentage of reads carrying deletions in TLC-CLIP and public CLIP libraries. **f** | Percentage of reads carrying insertion in TLC-CLIP and public CLIP libraries.

**Supplementary Fig. 8.**
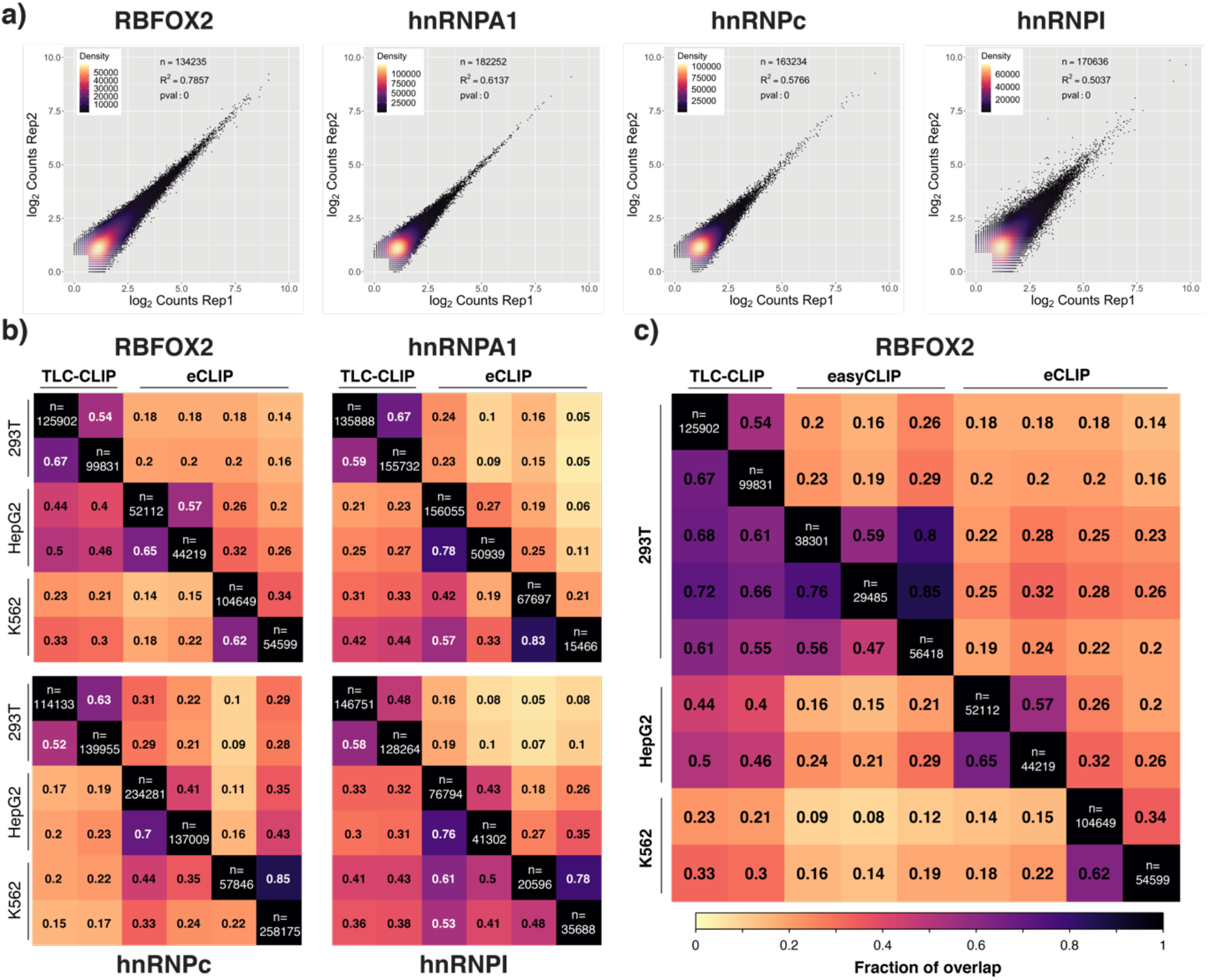
TLC-CLIP libraries are highly correlated and show good agreement with publicly available datasets at the peak-level. **a** | Read density plots displaying normalised log2 counts per TLC-CLIP replicate across concatenated peak sets for different RBPs. Correlation was calculated using Pearson correlation on log2 transformed normalised counts. **b** | Fraction of overlap at the peak level for different RBPs comparing TLC-CLIP with eCLIP libraries, requiring a minimum overlap of 25%between peaks. **c** | Fraction of overlap for RBFOX2 peaks (minimum overlap 25%) between TLC-CLIP, easyCLIP and eCLIP libraries, showing a strong agreement between TLC-CLIP and easyCLIP which were both generated in 293T cells.

**Supplementary Fig. 9.**
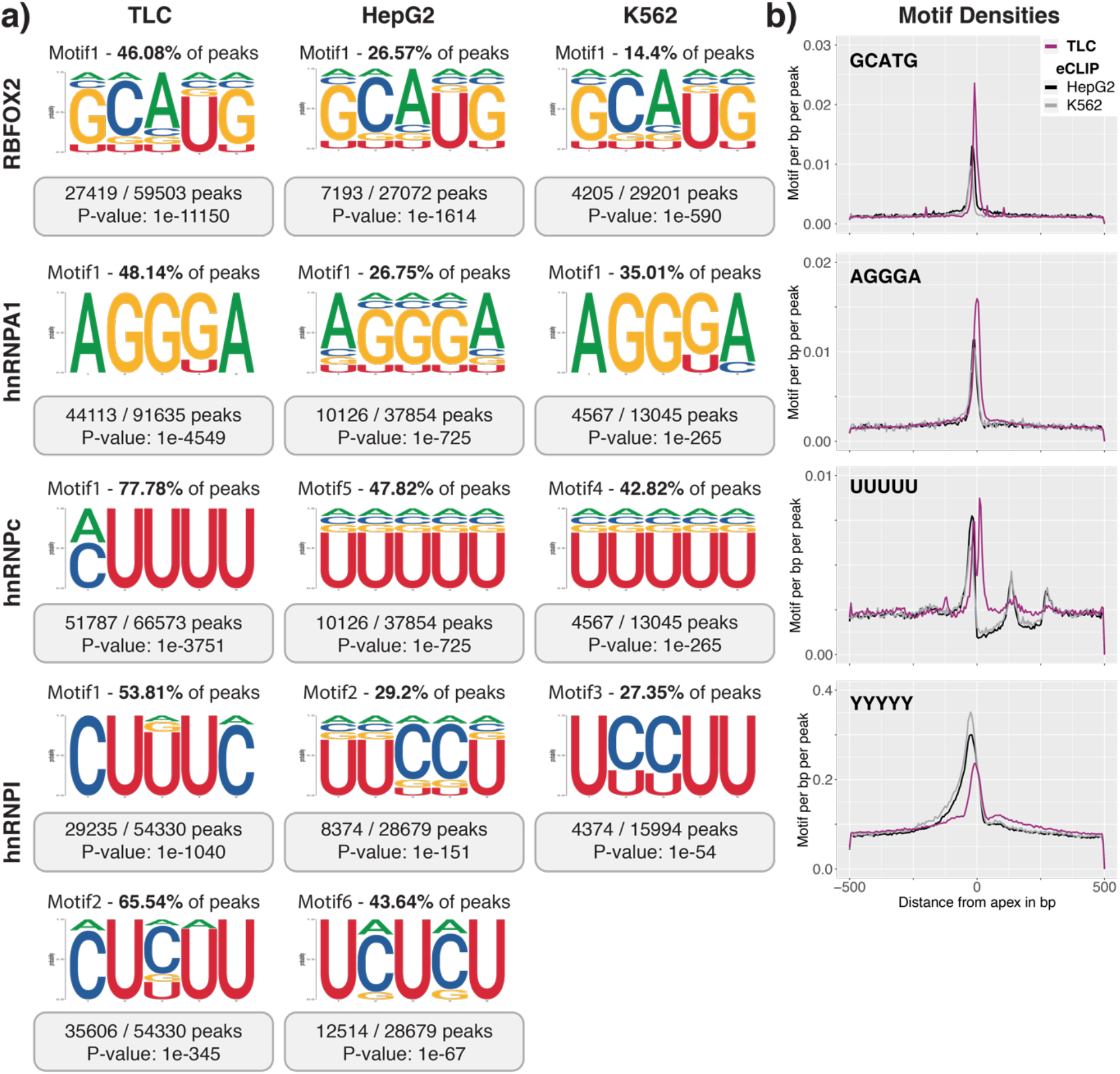
TLC-CLIP libraries allow *de novo* motif discovery and show strong motif enrichment. **a** | Position weight matrix from *de novo* motif discovery matching the known binding motif. Motif score, percentage of peaks carrying motif, as well as P-value obtained from the homer motif discovery software are stated. **b** | Normalised motif density plot displaying motifs per bp per peak for consensus motif across TLC-CLIP and eCLIP peaks, centred on the peak apex defined by CLIPper as the most likely site of crosslink.

**Supplementary Fig. 10.**
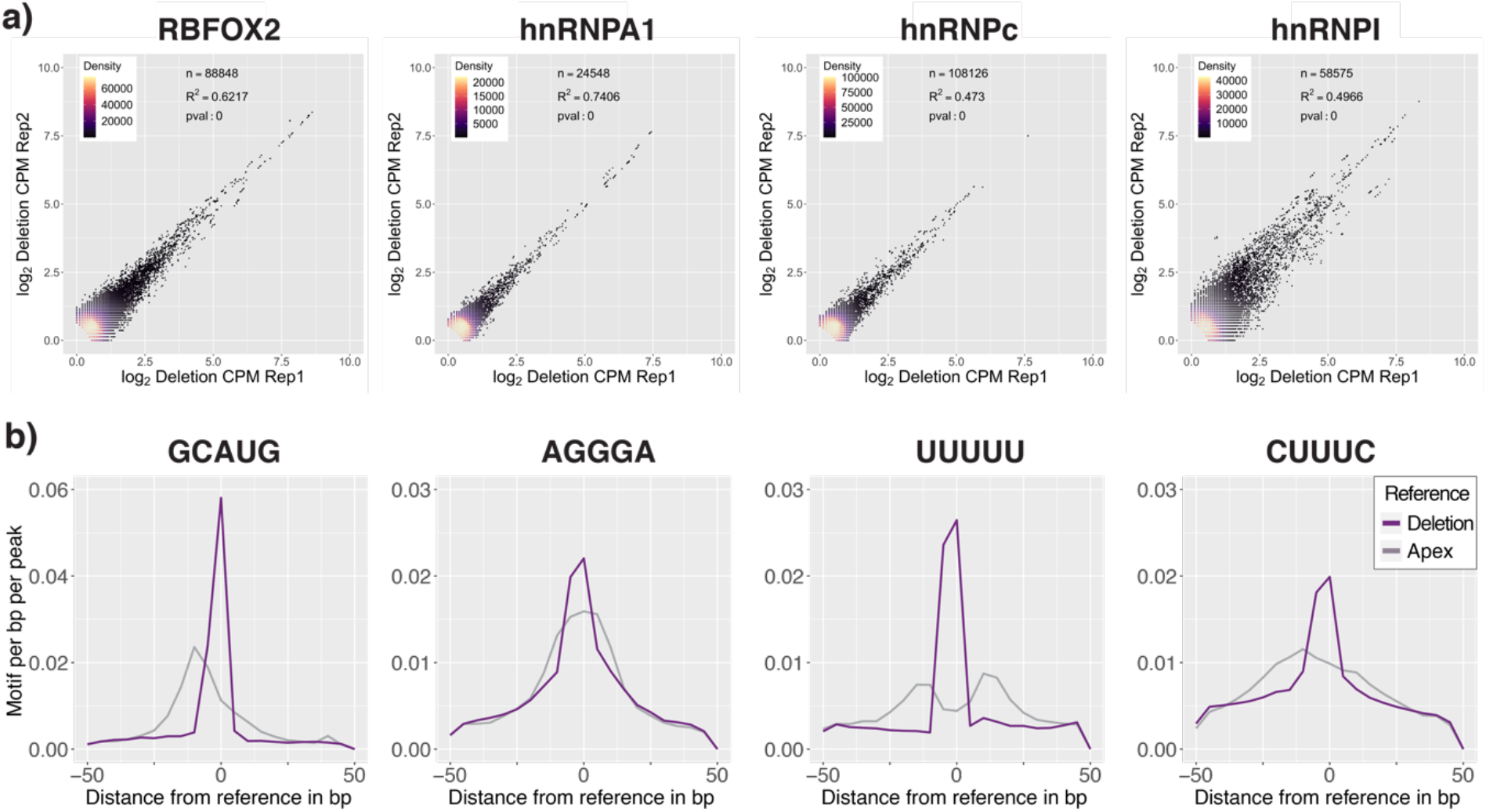
Crosslinking-induced deletions (CIDs) further increase the resolution of TLC-CLIP. **a** | Density plots displaying normalised log_2_ counts of crosslink-induced deletions per TLC-CLIP replicate at the single nucleotide level. Correlation was calculated using Pearson correlation on log2 transformed normalised counts. **b** | Motif density of consensus motif is increased when centring peaks on the position with the highest count of deletions compared with the apex region obtained from CLIPper.

**Supplementary Fig. 11.**
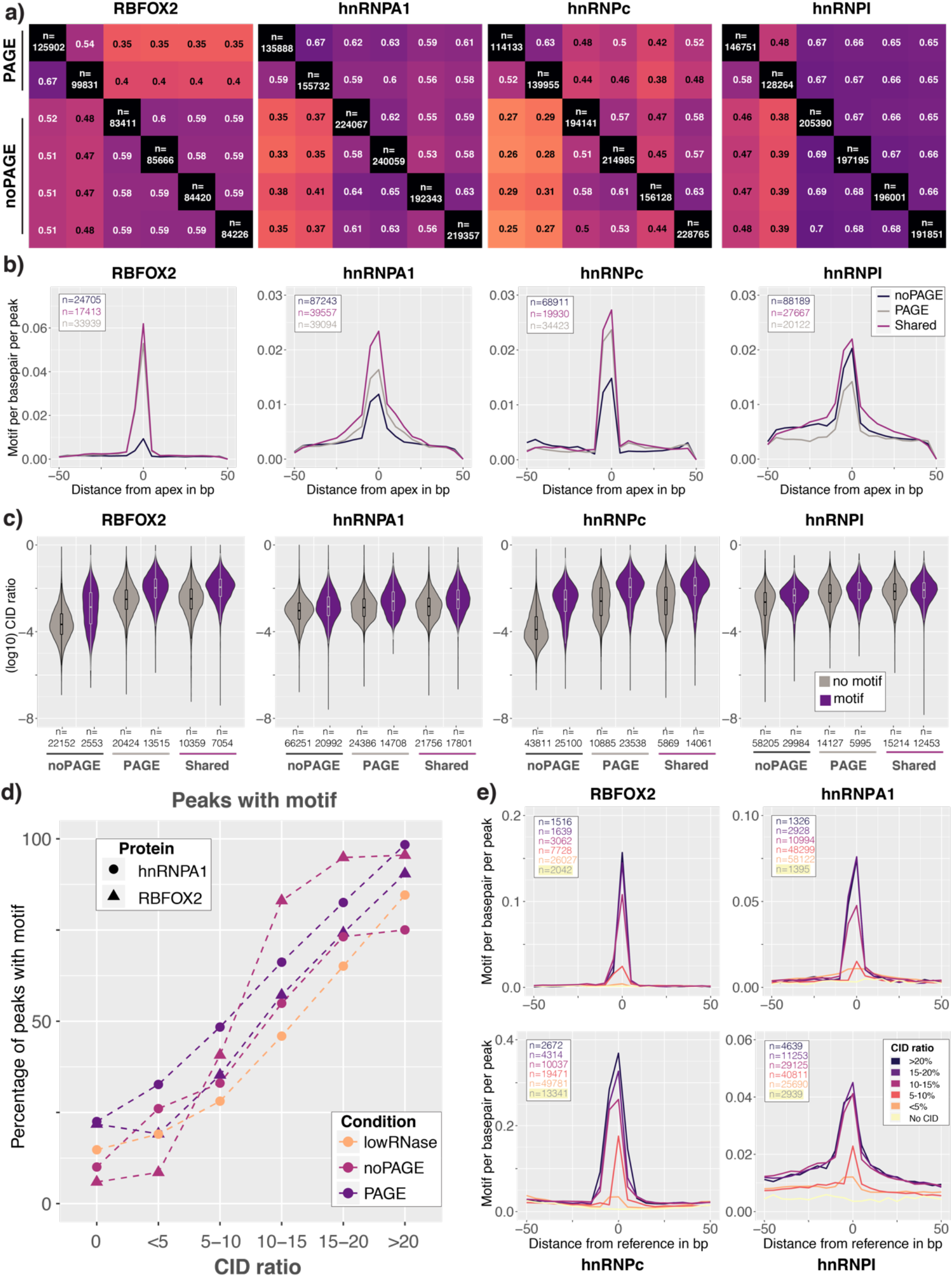
CIDs provide additional quality filter to increase specificity of TLC-CLIP libraries prepared without PAGE purification. **a** | Fraction of overlap between individual peaks obtained from TLC-CLIP libraries generated with or without PAGE purification, requiring a minimum overlap of 25%. For downstream analysis, peaks present in both PAGE replicates or in at least 3 out of 4 noPAGE replicates were used. **b** | Motif density of consensus motif across deletion centred peaks that are shared between experimental conditions or specific for libraries generated with or without PAGE purification **c** | Violin plots displaying the log_10_ of CID ratios for peaks grouped by experimental condition and annotated based on presence or absence of consensus motif. **d** | Percentage of peaks carrying consensus motif in CID peak subsets for different experimental conditions. **e** | Motif density of consensus motif across peak subsets according to their CID ratio.

**Supplementary Fig. 12.**
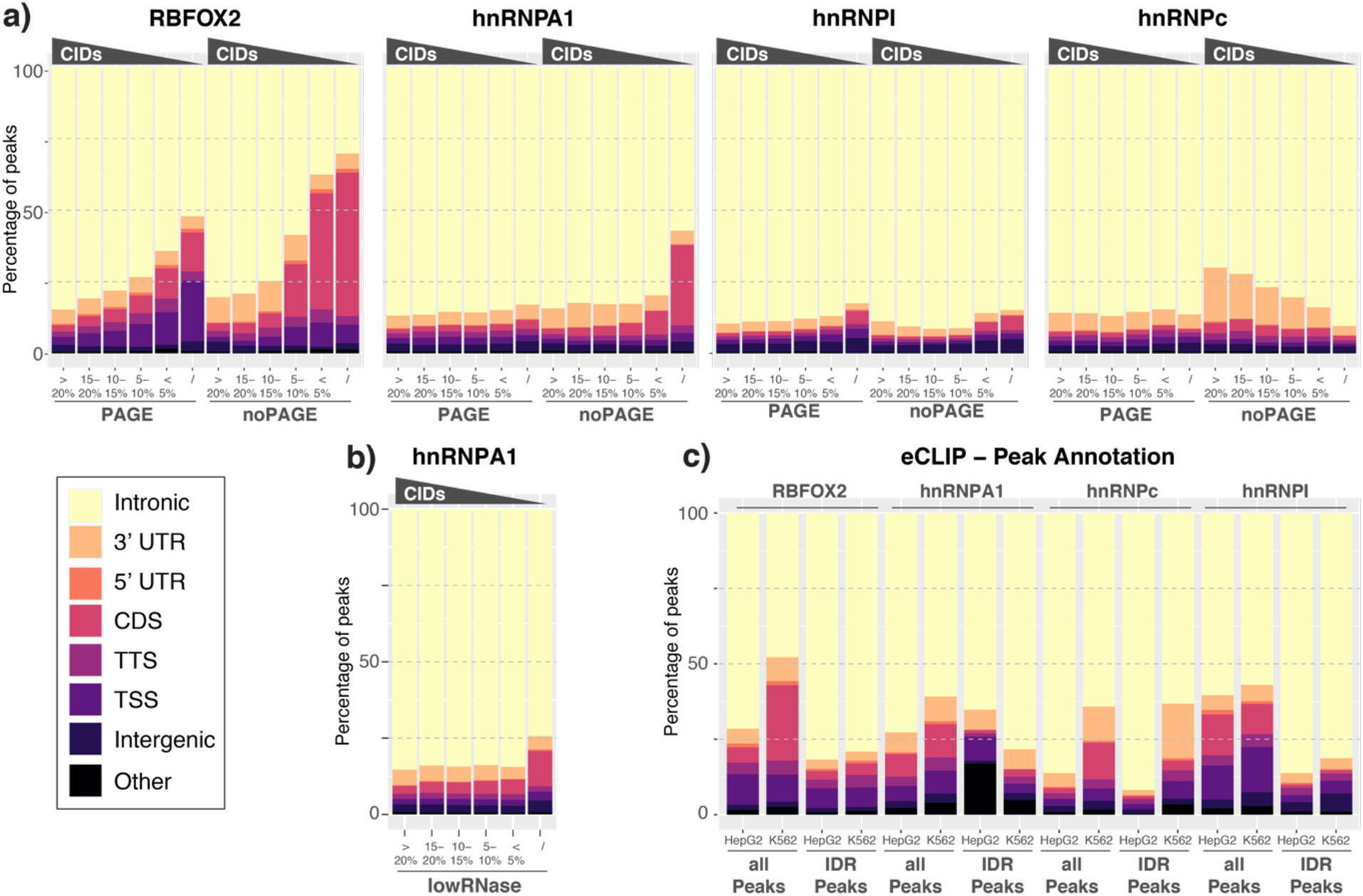
Peak annotation shows higher level of contaminating coding sequences among peaks with lower CID ratios. **a** | Percentage of peaks in TLC-CLIP libraries with varying CID ratios overlapping different genomic annotation layers (Intronic, 3’ UTR = 3’ untranslated region, 5’ UTR = 5’ untranslated region, CDS = coding sequences, TTS = −100 to +1kb around Transcription Termination site, TSS = -1kb - 100bp around Transcription Start Site; Intergenic, Other = including microRNA, non-coding RNA, pseudogenes, snoRNA and scRNA). **b** | Same annotation for hnRNPA1 libraries generated with low RNase condition (0.005U). **c** | Percentage of peaks in eCLIP libraries overlapping different annotation layers. Annotation was performed on CLIPper peaks with poisson cutoff of 0.01 and minimum score of 50 that are common between the two replicates per cell line (all Peaks) or on peaks filtered by Irreproducible Discovery Rate (IDR) obtained from encode.com which were first ranked by fold-enrichment over size-matched input controls (IDR peaks).

